# Pitfalls and recommendations for large-scale SNP genotyping in a non-model endangered species: the southern damselflies (*Coenagrion mercuriale*) as a case study

**DOI:** 10.1101/2024.05.02.592185

**Authors:** Agathe Lévêque, Jean-François Arnaud, Vincent Vignon, Clément Mazoyer, Cécile Godé, Anne Duputié

**Author notes:** Corresponding author: Anne Duputié.

## Abstract

Genomic markers are essential tools for studying species of conservation concern, yet non-model species often lack a genome reference. Here we describe a methodology for identifying and genotyping thousands of SNP loci in the southern damselfly (*Coenagrion mercuriale*), a bioindicator of freshwater stream quality classified as near-threatened, with locally declining populations. We used a hybrid approach combining reduced representation sequencing and target enrichment. First, we identified putative SNP loci using ddRADseq and *de novo* assembly. Then, single primer enrichment technology targeted 6,000 of these SNPs across 1,920 individuals. Challenges encountered included sequence recapture failure, coverage depth discrepancies, and aberrant *F*_IS_ values. We provide recommendations to address such issues. After multiple filtering, we retained 2,092 SNPs. We used them to characterise rear-edge populations of the southern damselfly in Northern France, a region where populations are sparsely distributed. Previous surveys utilising microsatellite markers allowed comparison of genetic diversity and differentiation estimates. Consistent with prior findings, genetic diversity estimates were similar across the studied populations that showed no sign of inbreeding. SNP markers exhibited greater resolution in detecting fine-scaled genetic structure, identifying two putative hybrids in adjacent populations, a feat unattainable with microsatellite loci. Altogether, this study highlighted the ongoing challenge of large-scale SNP genotyping using target sequencing techniques in non-model species to set conservation guidelines. Nonetheless, these new markers showed greater statistical power in identifying conservation units and offered the promise of greater precision in the identification of admixture events or the estimation of key population parameters such as effective population size.

## Introduction

Neutral molecular genetic markers are a crucial source of information in the fields of population genetics, evolutionary biology, and species conservation. They can be used for a wide range of applications, such as measuring the strength of micro-evolutionary processes of gene flow, genetic drift, mutation and selection, studying the spatial distribution of genetic diversity, estimating effective population sizes or delimiting conservation units. Therefore, it makes them essential tools in the establishment of conservation efforts, notably for endangered species (Allendorf *et al*., 2010; Holderegger *et al*., 2019; Hohenlohe *et al*., 2021; Allendorf *et al*., 2022). Many different approaches have been developed to genotype sets of molecular markers in a context of conservation biology, each with its own benefits and limitations (Davey *et al*., 2011; Allendorf, 2017). Because of their polymorphism, reproducibility, ease of development and genotyping, microsatellite markers were widely used to study natural populations (Frankham *et al*., 2013; Allendorf, Funk, Aitken, Byrne, Luikart, *et al*., 2022). However, in the last decade, these markers have rapidly been replaced by single nucleotide polymorphism (SNP) markers, which are more abundant and widely distributed across the genome, offering the promise of greater precision and statistical power in the estimation of population genetics parameters (Allendorf *et al*., 2010; Harrisson *et al*., 2014). The success of this new type of molecular marker is intimately linked to the development of next-generation sequencing (NGS) technologies, which has led to an increasingly large amount of genomic data being generated for wild populations (Harrisson *et al*., 2014; Allendorf, 2017; Hohenlohe *et al*., 2021). However, integrating these new genomic tools in conservation practices remains challenging (Allendorf *et al*., 2010; Shafer *et al*., 2015; Allendorf, 2017). Studying the spatial genetic structure of wild populations across space requires sampling hundreds or even thousands of individuals in multiple populations (Narum *et al*., 2013). Although it is now possible to sequence whole genomes for a subset of individuals, reduced genomic data can provide sufficient and valuable information to answer many scientific questions in conservation biology. Moreover, obtaining complete genome sequences for many individuals considerably increases the cost and complexity of bioinformatics analyses, particularly when the genome sequence of the focal species is unknown (Narum *et al*., 2013). Consequently, one of the main impediments to the application of genomic tools to the conservation of wild populations is the need to genotype thousands of individuals with thousands of genetic markers in species for which often, little or no reference genomic data is available (Seeb *et al*., 2011; Narum *et al*., 2013).

Various strategies for sequencing and genotyping a sub-part of the genome emerged as powerful alternatives to whole genome sequencing (Davey *et al*., 2011; Jones and Good, 2016). Among these approaches, reduced representation sequencing (RRS, Narum *et al*., 2013; Andrews *et al*., 2016; Campbell *et al*., 2018) and target enrichment methods (Mamanova *et al*., 2010) appeared to be complementary in terms of genomic knowledge required for their application, the effort and time required for their development, and their ability to genotype a large number of samples.

RRS approaches such as RAD-seq (Baird *et al*., 2008), genotyping-by-sequencing (GBS, Elshire *et al*., 2011) and double-digest RAD sequencing (ddRAD; Peterson *et al*., 2012) make it possible to identify and sequence many SNP markers in non-model species (Narum *et al*., 2013). These methods rely on the reduction of genomic complexity by using restriction enzymes to produce short-sequenced fragments that provide the frame for the discovery of SNPs widely spread across the genome (Van Tassell *et al*., 2008). The major advantage of these techniques is that they do not require any prior genomic information, which makes them particularly applicable in the conservation of non-model species for which there is often no available reference genome (Peterson *et al*., 2012). In addition, the development of ddRAD sequencing and associated computer tools (e.g. STACKS; Catchen *et al*., 2013) has made it possible not only to discover polymorphic sites but also to construct *de novo* consensus sequences (RADloci). However, these methods have some limitations, especially because the random distribution of enzymatic cut sites impedes targeting specific loci, which reduces the reproducibility between studies and spreads the sequencing effort over a very large number of sites (Andrews *et al*., 2016; Barchi *et al*., 2019; Scaglione *et al*., 2019).

In parallel with the development of RRS techniques, target enrichment methods arose as viable alternatives for obtaining cost-effective genome-wide genotypic data with higher reproducibility (Meek and Larson, 2019). These methods are based on the design of DNA capture beads or PCR amplification probes that target specific genomic regions, focusing sequencing efforts on the targeted regions, reducing the cost per sample, and thereby increasing the number of samples that can be genotyped (Mamanova *et al*., 2010; Jones and Good, 2016; Meek and Larson, 2019). Tecan Genomics (Redwood City, CA) developed the Allegro Targeted Genotyping system (ATG), a ready-to-use technique for custom targeting of previously identified SNPs (Scaglione *et al*., 2019). This technique presents the advantages of moderate cost and subcontracted probe development, whilst having a flexible design and ready-to-use library preparation kits. In addition, the use of individual combinations of barcodes allows pooling several thousand samples, and interrogating thousands of SNPs in a single sequencing run. Based on a single primer enrichment technology (SPET), ATG relies on the design of ∼40 bp long probes based on reference genomes or transcriptomes, enabling to target with high reproducibility specific genomic regions carrying genetic variants of interest. The method has so far mainly been used in human health (Scolnick *et al*., 2015; Nairismägi *et al*., 2016; Saber *et al*., 2017) or plants of agronomic interest (Barchi *et al*., 2019; Scaglione *et al*., 2019; Baccichet *et al*., 2022; Tripodi *et al*., 2023) and its use on wildlife species for conservation purposes is only starting to develop (Gramazio *et al*., 2020; Gavriliuc *et al*., 2022).

The main limit to the application of target enrichment methods is the need to have prior knowledge of the genome and the genetic variants to be targeted. To overcome this issue, several hybrid approaches were developed in recent years to genotype genomic variants in a very large number of samples at a reduced cost and without any prior genomic knowledge (Campbell *et al*., 2015; Ali *et al*., 2016; Meek and Larson, 2019). These methods combine a RRS approach to identify putative SNPs, which are then targeted for enrichment. Here, we applied such a hybrid method to develop and genotype thousands of SNPs in numerous samples of a non-model species with high conservation concerns, the southern damselfly *Coenagrion mercuriale* (Odonata, Zygoptera).

The southern damselfly is a particularly interesting study model to integrate evolutionary genomics with ecology, as it exhibits a complex biphasic (aquatic and terrestrial) life-cycle and represents a bioindicator of freshwater stream quality (Bybee *et al*., 2016; Watts *et al*., 2004c). The species is restricted to western and southern Europe, and became almost extinct in seven European countries on the eastern edge of its geographic distribution, where many populations are threatened by habitat loss or deterioration (Grand, 1996). Consequently, the southern damselfly is regarded as Near Threatened by the IUCN (Boudot, 2020), listed on Appendix II of the Bern Convention and on Annex II of the EC Habitat directive, in addition to a national protection in France where the southern damselfly is widely distributed and locally abundant, except in the northernmost regions (Houard, 2020; Fierimonte and Vanappelghem, 2021). In these northernmost regions, southern damselfly populations are stable but sparse, mainly due to modern intensive agricultural practices with the shift from grasslands to irrigated ploughed fields, in addition to the closure of waterways by scrubs and forests (Fierimonte and Vanappelghem, 2021). In northern France, southern damselfly populations are monitored by biodiversity managers, and major demographic declines were recorded with only a few individuals found in some years (Vanappelghem and Hubert, 2010). In this context, the conservation status of several populations in the region was previously described in the light of levels of genetic diversity and estimates of gene flow among populations (Lorenzo-Carballa *et al*., 2015; Lévêque *et al.,* 2024).

In this study, we used a two-step approach for the massive genotyping of thousands of SNPs in numerous southern damselfly populations. First, we used ddRADseq library preparation and a *de novo* assembly analysis to construct reference contigs and to identify SNPs markers using paired-end sequencing (Peterson *et al*., 2012; Rochette *et al*., 2019). Secondly, we used a single primer enrichment technology (Allegro Targeted Genotyping) to target six thousand SNPs and massively genotype thousands of southern damselfly individuals from various populations. We discuss the technical issues associated with the application of targeted enrichment genomic techniques on a non-model species. Finally, we carried out population genetic analyses to assess the levels of genetic diversity and of genetic differentiation, and to identify population genetic affiliation in a set of five threatened southern damselfly populations located in northern France. These populations were previously studied using microsatellite markers (Lorenzo-Carballa *et al*., 2015; Lévêque *et al.,* 2024), which enables us to compare our results with previous findings and to suggest monitoring actions of these populations.

## Materials and methods

### 1/ Reduced-representation genome sequencing and de novo SNP discovery

#### Sampling and DNA extraction

To build reference contig sequences and to identify SNPs present in diverse populations, we collected twenty males of southern damselflies from various locations in France in spring 2020 and 2021(see Table S1). Whole individual samples were stored in 100% ethanol prior to DNA extraction. Abdomens were removed to avoid sampling intestinal microbiota. We then crushed the remaining tissues using five MN Beads Type D (Macherey-Nagel, Duren, Germany). We extracted total genomic DNA from each sample using NucleoMag® Tissue Kit (Macherey-Nagel) according to the manufacturer’s recommendations. We then quantified DNA using a Qubit flex fluorimeter (Thermo Fisher Scientific, Waltham, Massachusetts, USA).

#### ddRAD library preparation and sequencing

To reduce the genome complexity, we prepared a double-digest restriction site-associated DNA sequencing (ddRADseq) library following the standard protocol described in Peterson *et al* (2012). In brief, 250 ng of DNA was digested with the restriction enzymes PstI (recognising the rare CTGCA/G motif) and MspI (recognising the frequent motif C/CGG; both enzymes from New England Biolabs, Ipswich, MA, USA). After the ligation of two adapters P1 and P2, each containing a unique barcode sequence, DNA was cleaned with 1X AMPure XP magnetic beads (Beckman-Coulter, Brea, CA, USA). Then, ligated fragments were amplified for 16 cycles using KAPA Hifi Hot Start PCR kit (Roche, Basel, Switzerland), and purified again with 1X AMPure XP beads. During the PCR, Illumina flow cell annealing sequences and a second index barcode were appended to the sequences, giving each sample a unique combination of barcodes for downstream identification. Next, after a size selection by AMPure XP double SPRI (0.5X and 1X), libraries were pooled in equimolar concentrations. A final control step was performed with Agilent 2100 BioAnalyzer, and the pooled libraries were then sequenced for 250 bp paired-end on a NovaSEQ6000 instrument (LIGAN-PM sequencing platform at EGID, Lille, France).

#### Building reference sequences and identifying SNPs

We used STACKS software pipeline v2.60 (Catchen *et al*., 2013; Rochette *et al*., 2019) following the *de novo* analysis protocol described by Rivera-Colón and Catchen (2022) to assemble RADloci from restriction-digested short-read sequences and to identify SNPs (Figure 1). First, raw sequence reads were cleaned and demultiplexed using the *process_radtags* module with the options to remove reads with any uncalled base (-c), discard reads with low-quality scores (-q) and rescue barcodes (-r) applied with default values. Sequence quality assessment was then performed using the software FastQC (https://www.bioinformatics.babraham.ac.uk/projects/fastqc/). All cleaned sequences were then trimmed to 200 bp using Trimmomatic v1.3.1 (Bolger *et al*., 2014). The *denovo_map.pl* pipeline (*ustacks*, *cstacks*, *sstacks, tsv2bam, gstacks* and *populations* modules) was then used to call SNPs and build the RADtag catalog, with the number of mismatches allowed to merge into one locus as M=9 (*ustacks*) and the number of mismatches allowed among individuals when building catalogs to n=9 (*cstacks*). These parameters were chosen after extensive exploration of the parameter space and choice of parameters appropriate for this dataset, following the R80 methods proposed by Paris *et al*. (2017). Finally, loci were filtered using the STACKS *populations* module to retain only loci shared by 80% of individuals (r = 0.8; Figure 1).

**Figure 1:**
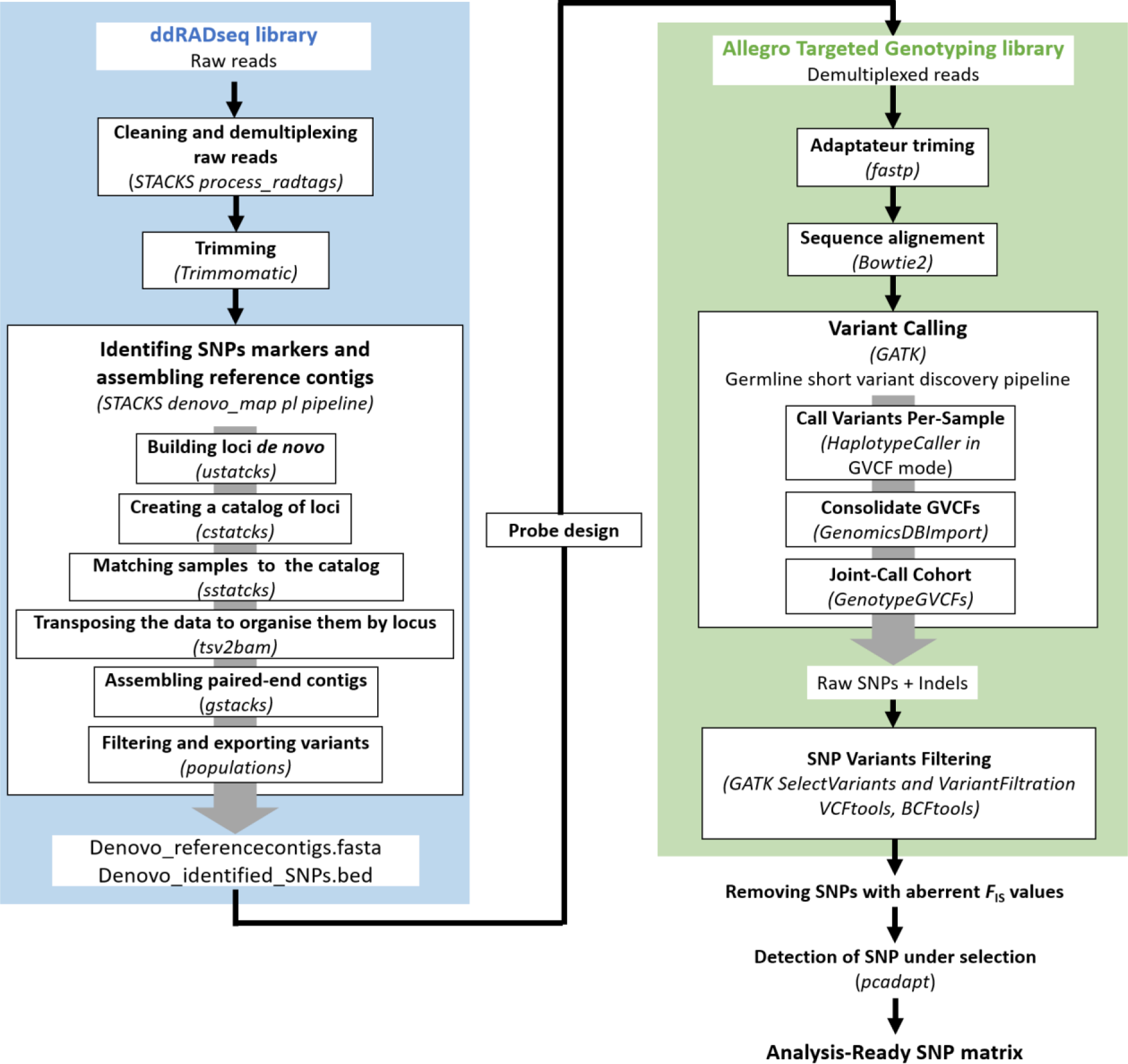
Bioinformatic pipeline for a two-step approach to massively genotype thousands of SNP markers in a non-model species, the southern damselfly (*Coenagrion mercuriale*). First, we used the STACKS software to identify *de novo* SNP variants and construct RADloci as reference sequences from a ddRADseq library (blue zone). Then, we massively genotyped the targeted SNPs through Allegro Targeted Genotyping developed by Tecan Genomics, and filtered the data to obtain high-quality genotypes (green zone).

### 2/ Massive SNP genotyping with Single Primer Enrichment Technology

#### Designing probes for target SNP loci

The Allegro Target Genotyping (Tecan Genomics Inc., Redwood City, CA) used in our assay implements Single Primer Enrichment Technology (SPET), which involves hybridisation of custom-designed probes near target SNPs, followed by probe extension, the addition of sequencing adapters, and high-throughput Illumina sequencing. We first selected SNPs out of the 80 bp of the start or end of the RADcontigs sequence and removed RADcontigs with uncalled bases (N), leaving 22,224 eligible SNPs distributed over 10,419 RADlocus. Probes for the target SNPs were about 40 bp long and were custom-designed by Tecan using the previously identified RADloci. Two probes were designed per target SNP, one per DNA strand. From the first panel of probes build by TECAN, we selected probes with a unique alignment, we kept only one pair per RADlocus and we then randomly sub-selected 6,000 couples of probes (Figure S1). Our final panel thus contained 12,000 probes targeting 6,000 SNPs occurring each on a different RADcontig (Table S2).

#### Sampling and DNA Extraction

We sampled 1,920 southern damselflies from various locations in France, with 1 to 32 individuals sampled per population (mean 16.55 ± 10.07). We collected them with an insect net in late spring (May to July) from 2020 to 2022. Whole individuals were stored in 100% ethanol, and DNA was extracted following the procedure described above. In the present paper, we only focus on 50 individuals sampled in Northern France to analyse the levels of genetic diversity and the extent of genetic differentiation. Indeed, these populations are all located at the edge of the geographic distribution of the species (Figure 2A) and were previously characterised using a set of ten microsatellite markers. This allows us to compare the genetic structure observed using classical microsatellite with that observed with the newly developed SNP markers (Lévêque *et al.,* 2024).

**Figure 2:**
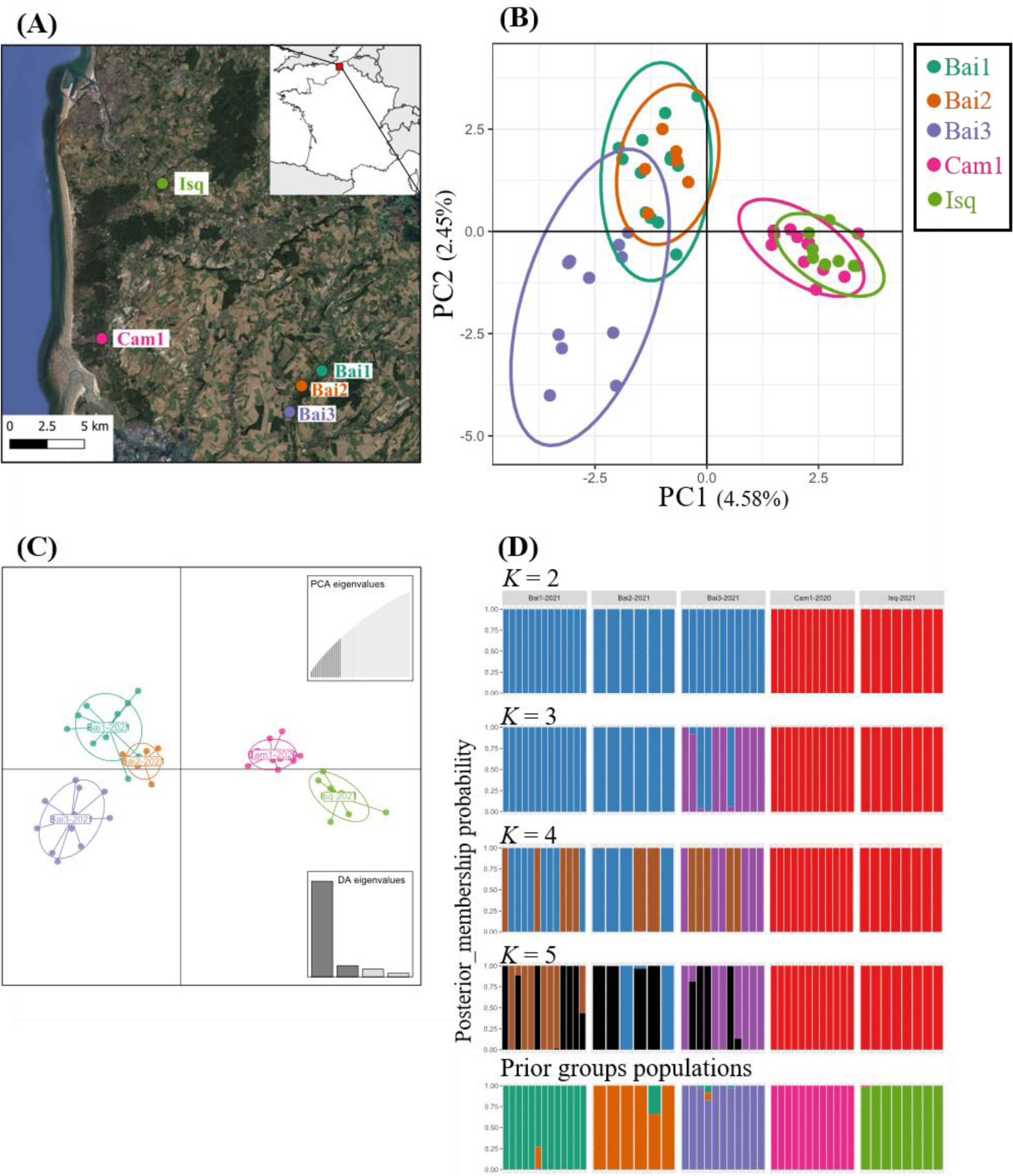
Patterns of genetic structure in five southern damselfly (*Coenagrion mercuriale*) populations located at range margin in northern France. **(A)** Map showing their geographical locations. **(B)** Principal Component Analysis (PCA), **(C)** Discriminant Analysis of Principal Components (DAPC) with sampling groups as priors, **(D)** assignment plots from DAPC for a number of clusters *K* varying from two to five with sampling groups as priors. Colors in panels B and C followed those in panel A; the same holds for panel D based on sampling populations as prior groups.

#### SPET library preparation

We followed the guidelines of the manufacturer available at https://www.tecan.com/hubfs/HubDB/Te-DocDB/pdf/UG_Allegro_Targeted_Genotyping_V2.pdf, to prepare the SPET libraries. We used the Allegro Metaplex Module, an optional module provided by TECAN to increase sample multiplexing capacity during library preparation. By default, Allegro Targeted Genotyping V2 includes 192 unique 1/i7 barcode for tagging samples. The Metaplex kit allows adding a second barcode (2/i5 barcodes) during the library amplification step. Sixteen 2/i5 barcodes are available from TECAN genomics: MP2-BC01 to MP2-BC16. We used the first ten 2/i5 barcodes, enabling all 1,920 samples to be identified in subsequent analyses by a unique combination of barcodes (Table S3). After qualitative and quantitative analysis of each pool of 48 individuals with a Qubit flex fluorimeter (Thermo Fisher Scientific, Waltham, Massachusetts, USA) and a 5200 Fragment Analyzer System (Agilent Technologies, Santa Clara, CA, USA), we pooled them equimolarly into a single tube containing all 1,920 samples. After a final quantity and quality control step was carried out, the pooled genomic library was sequenced on a MiSeq System (1x150 bp, Illumina Inc., San Diego, CA, USA sequenced at GenoScreen sequencing platform, Lille, France).

#### Sequence filtering and SNP genotyping

First, we removed low-quality reads and trimmed the Illumina adapters of the raw demultiplexed reads using *fastp* v0.23.4 (Chen *et al*., 2018) with the parameters -y, -5, -3 (enabling the low-complexity filter and dropping low-quality bases from both ends of the reads; see Figure 1). Then, we removed the 768 individuals that were tagged with i5 barcodes MP2-BC07 to MP2-BC10, because they had about 9 times fewer reads than those labelled with the six remaining i5 barcodes (Table S3, Table S4). We also removed all reads not starting exactly with a 40-base sequence corresponding to one of the probes (Table S4).

The remaining reads were then aligned to the previously built RADloci using *Bowtie2* v2.4.1 (Langmead and Salzberg, 2012), with local alignment and parameters -L 32 -D 20 -R 3 -N 0 -i S,1,3 (Figure 1). SNP calling was performed using *GATK*-4.0 v4.2.2.0 (DePristo *et al*., 2011) following the software best practices workflow for germline short variant discovery. Analyses included the following steps: (i) per-sample variant calling on target regions using *HaplotypeCaller* with default parameters to create a GVCF file for each sample; (ii) the GVCF files were then consolidated using GenomicsDBImport in order to improve scalability and speed the next step, (iii) joint genotyping using GenotypeGCVFs with default parameters to produce a set of joint-called variants; (iv) selection of SNPs using *SelectVariants* and quality filtering of SNPs using *VariantFiltration* (filter expression used : QD < 2, QUAL < 30, MQRankSum < -12.5, MQ < 30; Figure 1).

An extra filtration of the genotype matrix was performed using *VCFtools* v0.1.16 (Danecek *et al*., 2011) and *BCFtools* v1.12 (Danecek *et al*., 2021). We tested a range of filter thresholds at each filtering step: loci depth of coverage, amount of missing data per loci, minor allele frequency (MAF) and the number of missing data allowed per individual (Table S5). After evaluating the effect of the stringency of each of these filters, we chose to code genotypes with fewer than 8 reads in depth of coverage as missing, and to retain only biallelic SNPs present in >50% of individuals and with minor allele frequency >0.01. After that, we selected a single SNP per RADlocus (the one with the least missing data), thus reducing linkage disequilibrium issues in subsequent population genetic analyses (Waples *et al*., 2022). We then filtered out individuals with >40% missing SNPs. After that, we removed loci showing more than twice the mean read depth, because variants that have a high mean depth across all samples can be indicative of either repetitive regions or paralogs (Li, 2014; O’Leary *et al*., 2018;). Finally, we only kept SNPs showing *F*_IS_ values within a range of [-0.2 ;0.2]. This criterion aimed to exclude loci with excessive bias in homozygotes or heterozygotes, suggestive of biologically irrelevant genotypic structures (Figure 3). Indeed, surveyed populations were known to be under classical Hardy-Weinberg equilibrium using microsatellite data (Lorenzo et al. 2015; Lévêque *et al.,* 2024).

**Figure 3:**
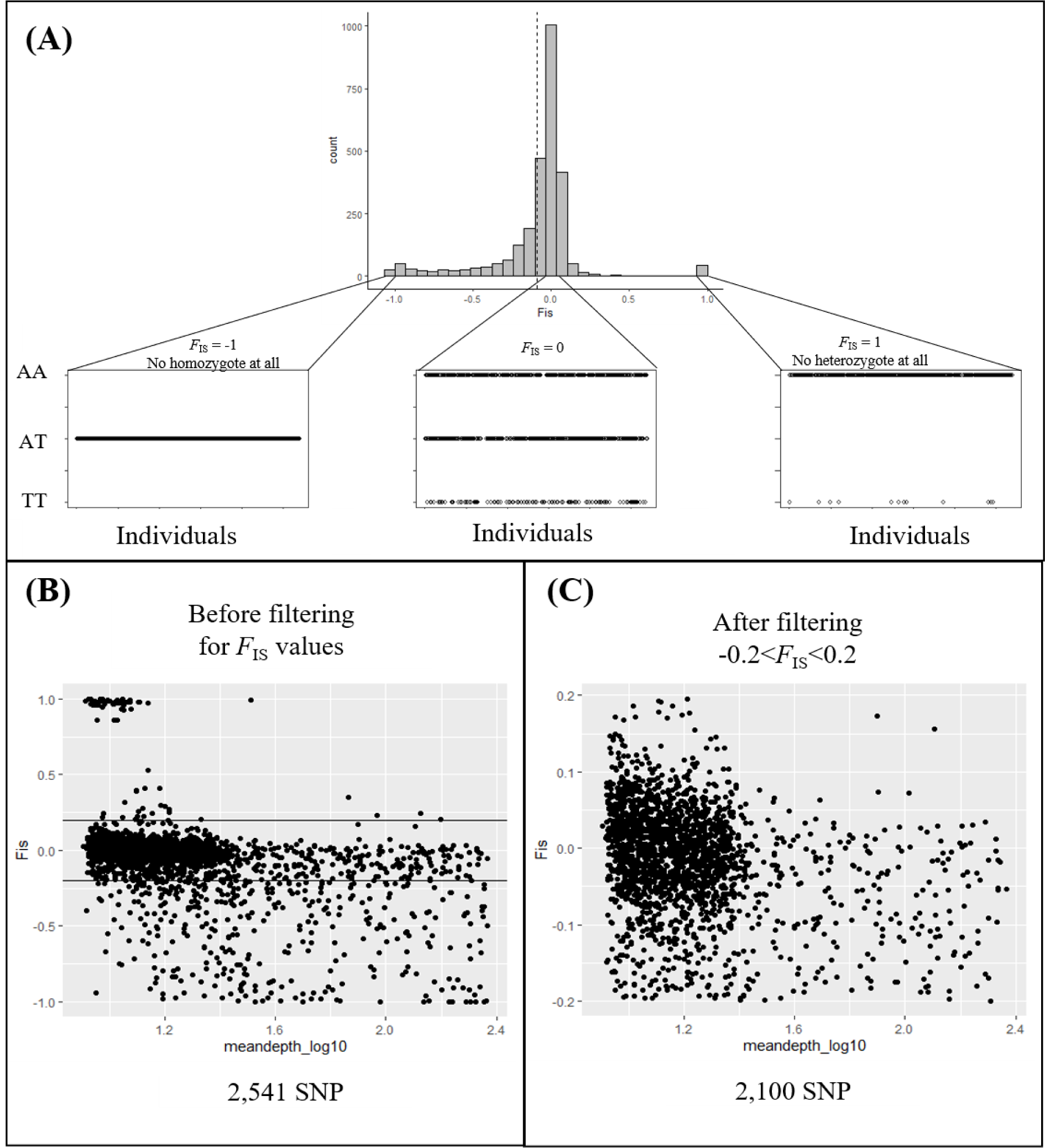
**(A)** Distribution of overall *F*_IS_ levels per locus in 1,080 individuals before filtering to retain only loci with *F*_IS_ levels between [-0.2 ; 0. 2], along with examples of genotypes of individuals for three loci characterised by *F*_IS_ =-1; *F*_IS_ =0, *F*_IS_ =1. **(B)** Before filtering on *F*_IS_ values, plot of the *F*_IS_ level per locus as a function of the average depth of reads on the RADloci carrying these SNPs, black bars indicate the limits of the filter that will be applied; **(C)** After filtering on *F*_IS_ values, plot of the *F*_IS_ level per locus as a function of the average depth of reads on the RADloci carrying these SNPs and distribution of *F*_IS_ levels per locus.

### 3/ Population genetic analyses

#### Genome scans for selection

We used the R v4.2.1 (R Core Team, 2022) package *pcadapt* v4.3.5 (Luu *et al*., 2017; Privé *et al*., 2020) to identify outlier loci potentially under selection, applying the protocol found at https://bcm-uga.github.io/pcadapt/articles/pcadapt.html, with the default Mahalanobis distance method. The number of principal components was selected using the scree plot method, which is based on Cattell’s rule stating to keep principal components up to the first inflexion point of the eigenvalues (Cattell, 1966). All SNPs showing *P*-values < 0.05 after adjustment for multiple testing through false discovery rate (Benjamini and Hochberg, 1995) were excluded from further analyses.

#### Genetic diversity, genetic differentiation, and population affiliation

We selected 50 individuals from five populations located in Northern France, towards the edge of the geographic distribution of the species. These populations were named Bai1, Bai2, Bai3, Cam1 and Isq and mapped on Figure 2A. The first three populations Bai1, Bai2 and Bai3 were located on the same watercourse and less than 2 km apart; populations Cam and Isq were more isolated and about 10-15 km away from populations Bai1-3.

We used the R packages *vcfR* v1.14.0 (Knaus and Grünwald, 2017), *dartR* v2.7.2 (Gruber *et al*., 2018; Mijangos *et al*., 2022), *poppr* v2.9.3 (Kamvar *et al*., 2015) to navigate between the file formats of the genotype matrix (vcf, genlight, genind). We estimated standard population genetic statistics (observed and expected heterozygosity, allelic richness) using the R library *hierfstat* v0.5-11 (Goudet, 2005). Allelic richness (*A*_r_) was rarefied on a minimal sample size of six individuals using the procedure of El Mousadik and Petit (1996).

To assess the levels of genetic differentiation between pairwise populations, we calculated pairwise *F*_ST_ estimates according to Weir and Cockerham (1984) using *StAMPP* v1.6.3 R package (Pembleton *et al*., 2013). Confidence intervals and significance of *F*_ST_ values were obtained using 1000 bootstraps over loci. Global levels of *F*_IS_ and *F*_ST_ were assessed using the R package *hierfstat* v0.5-11 (Goudet, 2005).

We then visualised the pattern of population affiliation using a principal component analysis (PCA) and a discriminant analysis of principal components (DAPC; Jombart *et al*., 2010), both implemented in the R package *adegenet* v2.1.9 (Jombart, 2008; Jombart and Ahmed, 2011). We used the function *find.cluster* for a number of clusters *K* varying from 1 to 5 to identify the optimal *K* (i.e. the one minimising the BIC). In addition, we also used the sampling populations as prior groups. Then, for groups built for *K*=2 to 5, we used a cross-validation procedure (*xvalDapc* with 1,000 replicates) to identify the optimal number of principal components retained to perform the DAPC, and subsequently applied it to assign individuals to a cluster and calculate their population membership probabilities.

## Results

### Reduced-representation genome sequencing and de novo SNP discovery

The Illumina NovaSeq 6000 SP 2x250bp sequencing produced a total of 1.28 billion reads, of which 1.12 billion (87.2%) passed initial quality filters and demultiplexing. Reads were filtered out due to ambiguous barcode sequences (6.56%), ambiguous RAD tags (6.18%) and low-quality scores (0.06%). After trimming reads to 200 bp and removing reads shorter than 200 bp, 199 million reads remained. The catalog generated by STACKS included 320,556 RADloci with a mean per-sample depth of coverage of 43.6X (s.d. 15.3X, from 19.6X to 82.3X). This number was reduced to 135,468 RADloci (mean length of 330.72 bp, s.e. 0.20) after requiring them to be present in at least 80% of the 20 genotyped individuals, resulting in a matrix of 758,786 SNPs.

### Massive SNP genotyping with Allegro Target Genotyping

The Illumina MiSeq SE 1x150bp sequencing produced a total of 1.784 billion single-end raw reads corresponding to an average of 929,223 reads per sample (from 54,628 to 3,926,551; Table S3). After adapter trimming and quality filtering, 99.98% of the initial reads remained. When removing the 768 individuals labelled with metaplexes MP2-BC07 to MP2-BC10 (which yielded too few reads per sample to be analysed; Table S3), and when removing sequences that did not begin exactly by a designed probe, the overall alignment rate on the RADloci was on average 99%. After SNP calling and the first quality filter, we identified 154,324 bi-allelic-SNPs distributed across 5,556 RADloci. Extra filtration of the SNPs produced a matrix of 2,541 SNPs containing 1,080 individuals genotyped for at least sixty percent of the loci (Table S5). After filtering out loci showing aberrant *F*_IS_ values, we obtained 2,100 high confidence SNPs (Figure 3). Eight SNPs were detected as outliers by *pcadapt* (keeping 2 PCs). These SNPs were therefore excluded to obtain a final dataset of 2,092 SNPs for population genetic analyses.

### Population genetic analyses

Using these newly developed SNPs, we observed similar levels of genetic diversity (*H*_o_, *H*_e_, *A*_r_) among the five surveyed populations located in Northern France (Table 1), with observed and expected heterozygosity ranging from 0.195 (populations Bai3 and Isq) to 0.207 (Cam1) and from 0.185 (Isq) to 0.205 (Cam1) respectively. Rarefied allelic richness ranged from 1.568 (Bai1) to 1.667 (Bai2) and *F*_IS_ levels ranged from -0.054 (population Isq) to -0.010 (for populations Bai2, Bai3, Cam1) (Table 1). As observed using these SNPs, microsatellite markers showed very similar patterns of genetic diversity among populations, with no evidence of departures from Hardy-Weinberg expectations (Table S6).

**Table 1:** Sampling information and mean multilocus estimates of genetic diversity using 2,092 biallelic SNPs in five southern damselfly (*Coenagrion mercuriale*) populations located at range margins of the species geographical distribution. Abbreviations: ID: population name. Longitude and latitude are provided (WGS84 ESPG 4326) coordinates. N: sample size; *H*_o_: observed heterozygosity; *H*_e_: expected heterozygosity; *F*_IS_: the intrapopulation fixation index; *A*_r_ allelic richness.

| ID | Sampling<br>Year | Longitude (E) | Latitude (N) | N | Allele<br>number | $H_o$ | $H_e$ | $F_{IS}$ | $A_r$ |
| --- | --- | --- | --- | --- | --- | --- | --- | --- | --- |
| Bai1 | 2021 | 1.8147 | 50.5564 | 13 | 3286 | 0.203 | 0.196 | -0.036 | 1.568 |
| Bai2 | 2021 | 1.7959 | 50.5472 | 6 | 2924 | 0.197 | 0.195 | -0.010 | 1.667 |
| Bai3 | 2021 | 1.7848 | 50.5314 | 11 | 3189 | 0.195 | 0.193 | -0.010 | 1.571 |
| Cam1 | 2020 | 1.6061 | 50.5738 | 12 | 3313 | 0.207 | 0.205 | -0.010 | 1.605 |
| Isq | 2021 | 1.6611 | 50.6682 | 8 | 2997 | 0.195 | 0.185 | -0.054 | 1.620 |

In terms of genetic differentiation, all populations showed significant levels of pairwise genetic differentiation, with *F*_ST_ values ranging from 0.014 to 0.108 (Table 2). Although geographically very close, the three populations located on the same watercourse (Bai1, Bai2, Bai3) showed significant pairwise *F*_ST_ ranging from 0.030 to 0.037 (Table 2). Compared with microsatellite data (mean multilocus *F*_ST_ = 0.068 and mean multilocus *F*_IS_=-0.089), the overall *F*_ST_ and *F*_IS_ in the studied region were of 0.055 and -0.022 respectively for the SNP data (Table S7).

**Table 2:** Pairwise *F*_ST_ estimates between the five southern damselfly (*Coenagrion mercuriale*) populations located in Northern France, associated with 95% confidence interval, and their statistical significance levels. * *P* ≤0.05, *** *P* ≤ 0.001.

|  | Bai1 | Bai2 | Bai3 | Cam1 |
| --- | --- | --- | --- | --- |
| Bai2 | 0.037<br>(0.023 - 0.051)*** |  |  |  |
| Bai3 | 0.030<br>(0.018 - 0.044)*** | 0.033<br>(0.019 - 0.049)*** |  |  |
| Cam1 | 0.051<br>(0.040 - 0.063)*** | 0.052<br>(0.037 - 0.068)*** | 0.076<br>(0.062 - 0.089)*** |  |
| Isq | 0.083<br>(0.067 - 0.099)*** | 0.080<br>(0.061 - 0.101)*** | 0.108<br>(0.088 - 0.127)*** | 0.014<br>(0.000 - 0.033)* |

With regards to population genetic affiliation, the PCA revealed a clear geographical and genetic distinctiveness between two sets of populations: (i) populations Bai1, Bai2, and Bai3, and (ii) populations Cam1 and Isq located further away (Figure 2B). For the DAPC, the optimal number of clusters was *K*=2, distinguishing the same two groups of populations (Figure 2D, Figure S3A). Visualising the DAPC results for *K*=3 clusters further allowed to separate specific individuals of populations Bai3 from those of populations Bai1 and Bai2 (Figure 2D, Figure S1B). With increasing numbers of clusters (*K*=4 and *K*=5), differences arose in individual assignment within populations Bai1, Bai2 and Bai3 (Figure 2D, Figure S1C,D). Finally, using sampled populations as a prior of assignation, most individuals were assigned to their own population, except two individuals who were assigned to two geographically close populations, suggesting admixture events between populations along the Bai river (Figure 2C,D).

## Discussion

### 1/ Pitfalls and recommendations when producing a SPET dataset for a non-model organism

Here we summarise and discuss the problems we faced, the solutions we explored, and the recommendations that emerged when genotyping thousands of individuals at thousands of SNPs loci using Allegro Targeted Genotyping (ATG) method, with contigs constructed from a ddRAD library for a non-model species with no genomic data available.

### Differences in ATG library DNA yields and their consequences on sequence data available

We used ten of the sixteen supplied Allegro i5 indexes (Table S3) to multiplex 1,920 samples using dual indexing of 48-sample pools. However, although the laboratory preparation conditions were identical for all pools, we obtained very different final DNA yields depending on the i5 indexes (Table S4). Indeed, those labelled MP2-BC07 to MP2-BC10 showed 2-3x lower DNA yields than the other six indexes. Even though the pools were brought back to equimolar proportions to form the final pooled library, we obtained around 9 times fewer reads for individuals labelled with these four indexes (Table S3).

As a consequence, all 768 individuals labelled with these four indexes exhibited low numbers of sequences resulting in a low SNP depth of coverage and subsequently a very low confidence in genotypes. Thus, we excluded all individuals marked with these four indexes from the rest of the analyses (resulting in only 1152 individuals being usable). Therefore, we recommend paying particular attention to the differences in the yield of Allegro libraries prepared with different indexes, because it may have a major impact on the resulting data, even if the pools are brought to equimolar concentrations prior to sequencing.

### Differential probe efficiency affects the number of suitable SNP

In Allegro Targeted Genotyping, the first 40 bp of each read should correspond to the sequences of a designed probe. We carried out preliminary analyses before the filtering, alignment and variant calling steps to check the sequences captured and the efficiency of the probes.

First, we examined whether all the sequences obtained began exactly with the 40 bp of the designed probes. 85.5% of the sequences we obtained did not exactly match any probe sequences, owing to insertions, deletions, or substitutions. This could indicate that during capture, probes may have hybridised with other non-targeted but genetically close portions of the genome. This may indicate possible paralogues or copy number variant in the southern damselfly genome (Verdu *et al*., 2016; Karunarathne *et al*., 2023). Consequently, we discarded sequences resulting from these likely parasitic hybridisations and whose first 40 base pairs did not exactly correspond to probe sequences.

Secondly, we checked which of the 12,000 designed probes were present in the first 40 base pairs of our sequences (0, 1 or 2 probes of the pair found, Table S4). Over the 6,000 initially designed pairs of probes, 704 were not found in any sequence. Consequently, the SNPs targeted by these probes cannot be found in our final dataset. Furthermore, on average over the 1,152 individuals remaining after the previous filtering step, only about half of the pairs of probes were completely recovered. Additionally, for about a third of the targeted SNPs, only one of the probes of the pair was found (Table S4). This drives variability in the sequencing depth between sites covered by two probes or by just a single probe (Scaglione *et al*., 2019; Gavriliuc *et al*., 2022).

### Aberrant departures from Hardy-Weinberg expectations

*F*_IS_ levels per locus were slightly negative on average, with many loci showing values close to zero, indicating panmixia. However, some loci showed extreme *F*_IS_ values, with *F*_IS_ = -1 (all individuals were heterozygous and no homozygote was found among all individuals) or *F*_IS_ = 1 (all individuals were homozygous but for different states, with no heterozygote occurrence; see Figure 3). Nonetheless, the same populations studied using microsatellite markers followed a Hardy-Weinberg equilibrium with non-significant *F*_IS_ levels (Lévêque *et al.,* 2024).

We hypothesised that extreme values in *F*_IS_ were due to differences in depth of coverage: strong excess in heterozygotes could be correlated with excessively high depths, indicating possible duplication of the target genome region, whereas biases toward excess in homozygotes could be due to too low sequencing depth, making it impossible to detect alternative variants (Song *et al*., 2016; Lou and Therkildsen, 2022; Karunarathne *et al*., 2023). However, there was no clear relationship between loci sequencing depth and the levels of *F*_IS_ (Figure 3), although loci with a high homozygote excess showed a slight tendency to exhibit low coverage.

We decided to filter our loci using *F*_IS_ bounds set to -0.2 and 0.2, to remove loci showing aberrant departures from Hardy-Weinberg equilibrium, given what is known on patterns of inbreeding in that species (Keller *et al*., 2012; Lévêque *et al.,* 2024; Watts *et al*., 2004a,b, Figure 2). From a strict population genetic affiliation point of view, these extreme *F*_IS_ values did not impact the delimitation of population boundaries, as the PCA and DAPC analyses carried out before and after the application of this filter on *F*_IS_ led to the same geographical patterns and similar conclusions in terms of genetic structuring whatever the threshold chosen (data not shown). Yet, such extreme values in *F*_IS_ can lead to highly false estimates of inbreeding levels and of effective population size (Hedrick, 2011; Allendorf, et al. 2022). We therefore strongly recommend that future users of SPET method check whether the *F*_IS_ values associated with each locus are biologically relevant, and verify any biases before carrying out any analyses devoted to describe intra-population genotypic structure before drawing any conclusions on mating system and on local level of inbreeding.

### 2/Population genetics of southern damselflies in Northern France

The southern damselfly populations located in northern France showed relatively similar levels of genetic diversity and genotypic structure (*H*_o_, *H*_e_, *A*_r,_ *F*_IS_, Table 1) as previously described using microsatellite markers (Lorenzo-Carballa *et al*., 2015). This pattern was still true in 2021 (Lévêque *et al.,* 2024, Table S6). Interestingly, all populations showed slightly negative *F*_IS_ values, indicating a slight excess of heterozygotes compared with what is expected according to the Hardy-Weinberg equilibrium, an imbalance that was already markedly observed with microsatellite markers (Table S6).

In contrast to what can be observed with microsatellite data, significant levels of genetic differentiation were observed between all pairs of populations in Northern France, with an overall *F*_ST_ value of 0.055, indicating that gene flow among populations may be spatially constrained. This was true even at short geographical distances below 2 km, among the three populations located along the Bai watercourse. Nonetheless, pairwise levels of genetic differentiation estimated with SNP markers were lower than those observed with microsatellite markers in the same region (see Table S7 and Lorenzo-Carballa *et al*., [2015]) and in more fragmented southern damselfly populations in the UK (Watts *et al*., 2005, 2004c).

Finally, the study of population genetic structure using PCA and DAPC assignments showed a clear distinctiveness between two groups (Bai river and more coastal populations), which makes sense from a geographical point of view. Even at a finer spatial scale of less than 4km, population Bai3 appeared to be distinct from populations Bai1 and Bai2. This fine-scale population genetic distinctiveness is consistent with individual assignments obtained using a Bayesian clustering approach conducted on a similar set of populations using microsatellite markers (Lorenzo-Carballa *et al*., 2015).

Thanks to these patterns of fine-scale spatial structure and genetic differentiation identified using SNP markers, the DAPC analysis identified two individuals as of mixed origin between populations Bai1 and Bai2, which may be considered as putative first-generation hybrid in each of these two populations. This is consistent with previous studies reporting that southern damselflies can migrate over short distances of about 2 km (Rouquette and Thompson, 2007; Lévêque *et al.,* 2024; Watts et al., 2004c). From a conservation point of view, this suggested that the connectivity of the population Bai3 belonging to the same watercourse population should be improved by habitat management.

More generally, the ecological methods and genetic tools used to study Odonata populations already provided relevant information on dispersal pathways and landscape barriers among populations (Watts *et al*., 2006, 2007; Keller *et al*., 2010; Van Strien *et al*., 2012). These studies also highlighted a loss of genetic diversity in geographically isolated populations. Yet, little information is available on the evolutionary consequences of genetic erosion in Odonates, including the adaptative processes involved. Genomic techniques offer the potential to address these questions (Bybee *et al*., 2016).

## Conclusion

We showed that the ddRADseq RRS approach allows to identify *de novo* hundreds of thousands of SNP markers in a non-model species of conservation concerns (Peterson *et al*., 2012; Narum *et al*., 2013). However, the target enrichment method, initially designed to target 6,000 SNPs in 1,920 individuals, only managed to recover 2,100 SNPs for 1,080 individuals. This low recapture rate sharply contrasted with rates achieved in species where reference genomic sequences were available for developing Allegro probes (Barchi *et al*., 2019; Scaglione *et al*., 2019; Tripodi *et al*., 2023). This highlighted the ongoing challenge of large-scale SNP genotyping using targeted sequencing techniques in a non-model species. Future work using a combination of genomic and transcriptomic data targeting gene duplication, DNA loss, intron size variation or transposable elements may explain the difficulties we encountered in this study, but also may elucidate the putative mechanisms responsible for genome size variation in odonate species (Bybee *et al*., 2016). However, these transcriptomic and genomic studies would benefit from the availability of transcriptomes and reference genomes that need to be developed in Odonates (Bybee *et al*., 2016; Ioannidis *et al*., 2017).

However, the set of SNPs markers we developed in this study provided valuable information on the levels of genetic diversity, genetic differentiation and the spatial genetic structure of populations in local populations initially thought to be genetically depauperate. Notably, these markers allowed the identification of fine-scale genetic structure and patterns of gene flow that were undetectable using microsatellite loci. These newly developed SNPs thus emerge as valuable resources for future conservation genetics studies designed to infer crucial population genetic parameters, such as effective population sizes and migration pathways in southern damselfly populations.

