## Supplementary Figures and Tables for "Pitfalls and recommendations for large-scale SNP genotyping in a non-model endangered species: the southern damselflies (*Coenagrion mercuriale*) as a case study"

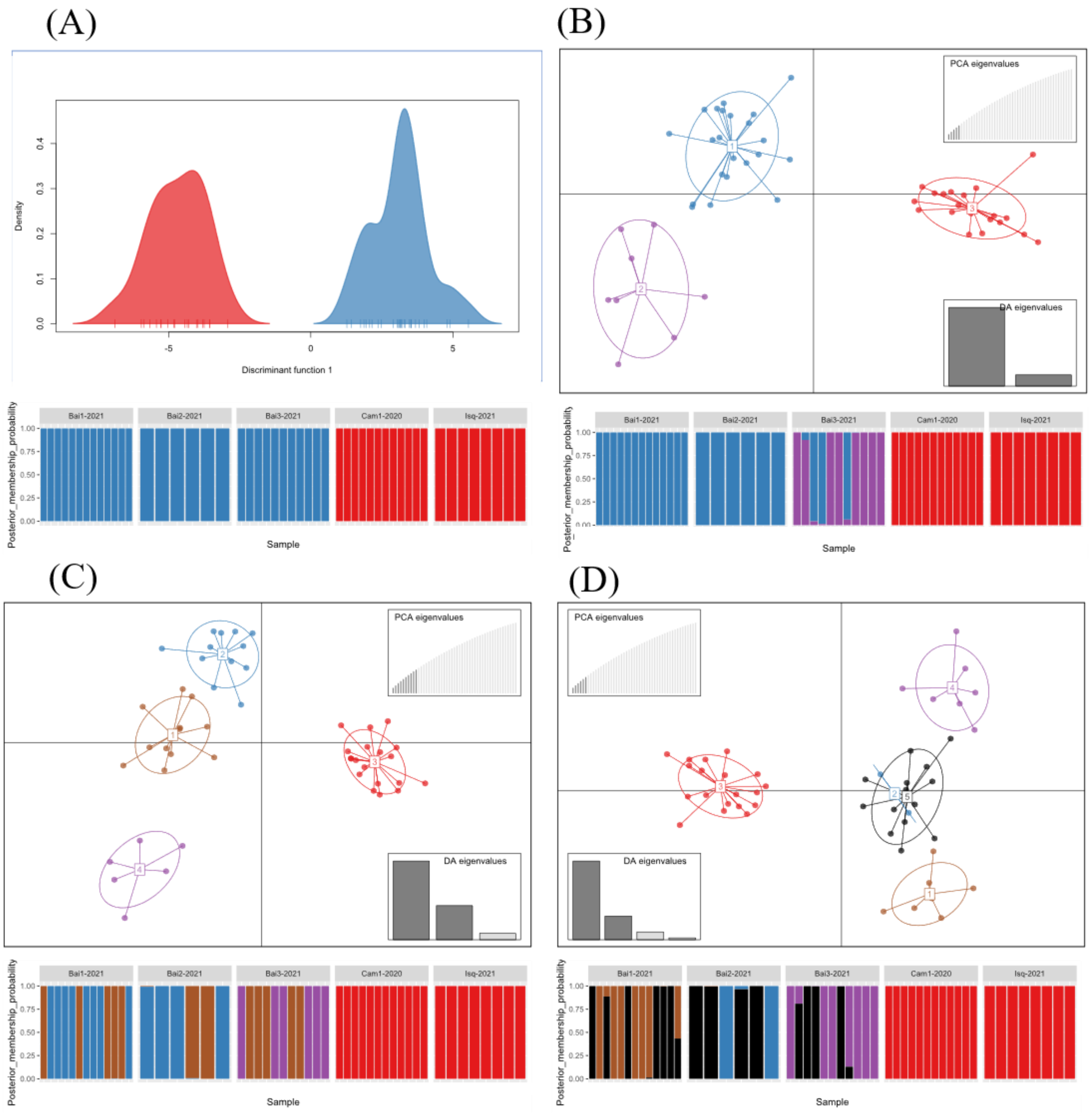


**Figure S1:** Scatterplots for DAPC for five southern damselfly (*Coenagrion mercuriale*) populations located in northern France with *K*=2 **(A)**, *K*=3 **(B)**, *K*=4 **(C)** and *K*=5 **(D)** with their associated population membership probability assignment.

**Supplementary Table S1:** List of the samples used in ddRADseq library preparation. Sample_ID: sample name, Pop_ID: name of the population of origin and geographic coordinates of the sampling sites (WGS84 ESPG 4326).

| Sample_ID | Pop_ID | Longitude (WGS84) | Latitude (WGS84) |
|---|---|---|---|
| Cam1-13-m20 | Cam | 1.60616400 | 50.57378824 |
| Isq-01-m21 | Isq | 1.66095648 | 50.66824458 |
| Bai1-01-m21 | Bai1 | 1.81473148 | 50.55641697 |
| Bai3-01-m21 | Bai3 | 1.78482142 | 50.53136707 |
| Cuy-01-m21 | Cuy | 3.45869472 | 50.44209135 |
| Ger1-05-m21 | Ger1 | 7.71086409 | 48.39779464 |
| Rut2-01-m21 | Rut2 | 7.59587037 | 48.49964518 |
| Ceh1-09-m21 | Ceh1 | 7.58164061 | 48.47838244 |
| S6-20-m21 | S6 | 7.61528337 | 48.62932940 |
| Rsc1-02-m21 | Rsc1 | 7.73923393 | 48.48230098 |
| A11-17-m21 | A11 | 7.65574505 | 48.57464250 |
| S21-06-m21 | S21 | 7.83357726 | 48.63317546 |
| Rla6-16-m21 | Rla6 | 7.70094941 | 48.67059395 |
| A5-08-m21 | A5 | 7.56451122 | 48.54390458 |
| V1-07-m21 | V1 | 7.60749656 | 48.19228039 |
| V2-15-m21 | V2 | 7.67439857 | 48.27871207 |
| RLK2-18-m21 | RLK2 | 7.65126778 | 48.65764007 |
| S17-10-m21 | S17 | 7.74115324 | 48.64159254 |
| V3-11-m21 | V3 | 7.73287300 | 48.35805528 |
| Ger6-14-m21 | Ger6 | 7.67472936 | 48.32892893 |

**Supplementary Table S2:** Details of the 12,000 SPET probes used, with their sequence (probe sequence), orientation (Strand) and the position on the reference RADlocus. The data table can be downloaded from the following link: https://figshare.com/s/a5aa49343eb1d9cd8093

**Supplementary Table S3:** i5 index, DNA pool and yields of the Allegro Target Genotyping library preparation.

| i5 index | Barcode Plate | Pool | Pool concentration (ng/µL) | Number of raw reads with this i5 index |
|---|---|---|---|---|
| MP2-BC01 | Plate 1 | 1 | 20.0 | 286 353 103 |
| | | 2 | 17.2 | |
| | Plate 2 | 3 | 14.2 | |
| | | 4 | 17.0 | |
| MP2-BC02 | Plate 1 | 5 | 15.9 | 265 877 177 |
| | | 6 | 13.9 | |
| | Plate 2 | 7 | 15.4 | |
| | | 8 | 12.5 | |
| MP2-BC03 | Plate 1 | 9 | 14.3 | 309 775 843 |
| | | 10 | 16.2 | |
| | Plate 2 | 11 | 20.6 | |
| | | 12 | 20.2 | |
| MP2-BC04 | Plate 1 | 13 | 13.2 | 207 371 842 |
| | | 14 | 9.72 | |
| | Plate 2 | 15 | 12.2 | |
| | | 16 | 9.42 | |
| MP2-BC05 | Plate 1 | 17 | 18.9 | 277 622 776 |
| | | 18 | 17.9 | |
| | Plate 2 | 19 | 19.1 | |
| | | 20 | 24.0 | |
| MP2-BC06 | Plate 1 | 21 | 22.8 | 313 291 482 |
| | | 22 | 18.4 | |
| | Plate 2 | 23 | 25.8 | |
| | | 24 | 22.6 | |
| MP2-BC07 | Plate 1 | 25 | 7.0 | 23 238 462 |
| | | 26 | 6.1 | |
| | Plate 2 | 27 | 7.3 | |
| | | 28 | 5.7 | |
| MP2-BC08 | Plate 1 | 29 | 7.5 | 31 631 178 |
| | | 30 | 6.2 | |
| | Plate 2 | 31 | 7.7 | |
| | | 32 | 6.4 | |
| MP2-BC09 | Plate 1 | 33 | 5.2 | 26 460 956 |
| | | 34 | 4.6 | |
| | Plate 2 | 35 | 6.0 | |
| | | 36 | 5.9 | |
| MP2-BC10 | Plate 1 | 37 | 6.8 | 42 484 721 |
| | | 38 | 6.4 | |
| | Plate 2 | 39 | 5.3 | |
| | | 40 | 5.0 | |

**Supplementary Table S4:** Information on southern damselflies samples used for the ATG. Sample ID, Year of sampling, Sampling site name and coordinates (WGS84), the metaplex used to tag the sample, the total and filtered number of sequences retrieved, the number of couples of probes completely (+ and - probe), partly (+ or - probe) or not at all found in the cleaned reads. Samples corresponding to metaplexes MP2-BC07, MP2-BC08, MP2-BC09 and MP2-BC10 which had to be withdrawn due to low numbers of reads, are shown in red. Northern France samples used for population genetic analyses in the present paper appear in bold face. The data table can be downloaded from the following link: https://figshare.com/s/a3514773186a615bbaed

**Supplementary Table S5.1:** Final steps for filtering the SNP matrix, showing the number of individuals and loci before and after each step. The final numbers of loci and individuals used for analysis are indicated in bold.

| **Filter** | **Before** | **After** |
|---|---|---|
| 1. Remove individuals tagged with i5 indexes MP2-BC07,08,09,10. | 1,920 individuals | 1,150 individuals |
| 2. Remove poor quality SNP (QD < 2, QUAL < 30, MQRankSum < -12.5, MQ < 30) | 254,174 SNPs | 174,822 SNPs |
| 3. Select only biallelic loci | 174,822 SNPs | 154,324 SNPs |
| 4. Select loci out of the 40bp corresponding to ATG probe sequences | 154,324 SNPs | 147,508 SNPs |
| 5. Set genotypes with depth <8x as missing data | 147,508 SNPs | 147,508 SNPs |
| 6. Discard loci genotyped in <50% of individuals | 147,508 SNPs | 58,053 SNPs |
| 7. Discard loci with minor allele frequency < 1% | 58,053 SNPs | 10,435 SNPs |
| 8. Select one loci per RADlocus (the one with the least missing data) | 10,435 SNPs | 2,767 SNPs |
| 9. Discard individuals with >40% missing loci | 1150 individuals | **1080 individuals** |
| 10. Remove loci with a max depth of coverage twice the mean coverage | 2,767 SNPs | 2,541 SNPs |
| 11. Filter out loci with $F_{IS}$ levels out of [-0.2;0.2] | 2,541 SNPs | 2,100 SNPs |
| 12. Filter putatively adaptative loci identified with pcadapt | 2,100 SNPs | **2,092 SNPs** |

We explored the effect of all these filtering parameters:

| | |
|---|---|
| TableS5.2 | Loci sequencing depth |
| TableS5.3 | Loci missing data amount |
| TableS5.4 | Minor Allele Frequency (MAF) |
| TableS5.5 | Individual missing data amount |

**Supplementary Table S5.2:** Effect of changing the filter for coverage depth of loci on the number of remaining RADloci (step 5 in Supplementary Table 5.1)

Firstly, we filtered our loci according to their sequencing depth (step 5 in table S5.1). We had two filtering options:

a) filter the loci according to their average depth of coverage (average per locus over all the individuals) with the risk of keeping loci with a fair average coverage but a low depth of coverage for some individuals.

b) Filter by locus and by individuals, by setting a threshold value above which we considered the genotype to be reliable, and transforming unreliable genotypes into missing data, with the risk of increasing the quantity of missing data in our dataset.

We elicited to choose the second option with a minimum coverage depth of 8X to consider genotypes as reliable.

| | **Depth** | **Number of biallelic SNP** | **Number RADcontigs containing biallelic SNP** | **Number of biallelic SNP located after position 40** | **Number RADcontigs containing biallelic SNP after position 40** |
|---|---|---|---|---|---|
| **Average depth of coverage filter** | 20X | 7002 | 830 | 6822 | 823 |
| | 15X | 14026 | 1468 | 13743 | 1466 |
| | 10X | 37489 | 2556 | 36451 | 2554 |
| | 8X | 60442 | 3208 | 58242 | 3205 |
| **Unreliable genotypes set as missing data** | 8X | 154324 | 5556 | 147508 | **5550** |

**Supplementary Table S5.3:** Effect of the filter on the amount of missing data per locus on the remaining number of RADcontigs (step 6 in Supplementary Table S5.1). We chose the last option.

| | Depth Filter | Loci present in x% of individuals | Number of biallelic SNP | nb RADcontigs containing biallelic SNP | Number of biallelic SNP after 40bp | Number RADcontigs containing biallelic SNP after 40bp |
|---|---|---|---|---|---|---|
| **Average depth of coverage** | **8X** | **90%** | 58950 | 3185 | 56797 | 3182 |
| | | **80%** | 59697 | 3204 | 57528 | 3201 |
| | | **70%** | 59870 | 3206 | 57694 | 3203 |
| | | **60%** | 59942 | 3206 | 57764 | 3203 |
| | | **50%** | 59988 | 3206 | 57807 | 3203 |
| **Unreliable genotypes set as missing data** | **8X** | **90%** | 12514 | 1419 | 12267 | 1415 |
| | | **80%** | 21362 | 1943 | 20917 | 1940 |
| | | **70%** | 32920 | 2415 | 32092 | 2411 |
| | | **60%** | 45910 | 2836 | 44442 | 2834 |
| | | **50%** | 60265 | 3198 | **58053** | **3195** |

**Supplementary Table S5.4:** Effect of the filter on Minor Allele Frequency (MAF) on the number of RADcontigs (step 7 in Supplementary Table S5.1). We chose the last option.

| Depth Filter | Locus present in x% of individual | Minor Allele Frequency | Number of biallelic SNP | Number RADcontigs containing biallelic SNP | Number of biallelic SNP after 40bp | Number RADcontigs containing biallelic SNP after 40bp |
|---|---|---|---|---|---|---|
| | | **<0.1** | 6560 | 2203 | 6465 | 2199 |
| **8X** | **50%** | **<0.05** | 7904 | 2451 | 7793 | 2444 |
| | | **<0.01** | 10605 | 2773 | **10435** | **2767** |

**Supplementary Table S5.5:** Effect of the filter on the amount of missing data in each individual on the number of RADcontigs (step 9 in Supplementary Table S5.1). We chose the third option.

| Filter individuals with less than X% SNP genotyped | Individuals kept after filtration. |
|---|---|
| **20%** | 1149 |
| **50%** | 1124 |
| **60%** | **1080** |
| **70%** | 965 |

**Table S6:** Sampling information and mean multilocus genetic diversity estimates using 10 microsatellite loci (Lévêque et al *biorxiv*) in five Southern damselfly populations. Abbreviations: ID: population name. Longitude and latitude are provided in WGS84 coordinates. *N*: sample size. $H_o$: observed heterozygosity. $H_e$: expected heterozygosity. $F_{IS}$: the intrapopulation fixation index; $A_r$ allelic richness.

| ID | Sampling Year | Longitude (E) | Latitude (N) | *N* | Allele number | $H_o$ | $H_e$ | $F_{IS}$ | $A_r$ |
|---|---|---|---|---|---|---|---|---|---|
| Bai1 | 2021 | 1.8147 | 50.5564 | 25 | 37 | 0.544 | 0.49 | -0.110 | 3.071 |
| Bai2 | 2021 | 1.7959 | 50.5472 | 11 | 30 | 0.518 | 0.484 | -0.070 | 2.866 |
| Bai3 | 2021 | 1.7848 | 50.5314 | 14 | 28 | 0.529 | 0.44 | -0.202 | 2.707 |
| Cam1 | 2020 | 1.6061 | 50.5738 | 21 | 38 | 0.51 | 0.489 | -0.043 | 3.229 |
| Isq | 2020 | 1.6611 | 50.6682 | 20 | 31 | 0.485 | 0.461 | -0.052 | 2.820 |

**Table S7:** Pairwise $F_{ST}$ estimates using 10 microsatellite loci (Lévêque et al *biorxiv*) between five southern damselfly populations located in Northern France, associated with their 95% confidence interval, and their statistical significance levels. * $P \leq 0.05$, ** $P \leq 0.01$ *** $P \leq 0.001$.

| | Bai1 | Bai2 | Bai3 | Cam1 |
|---|---|---|---|---|
| Bai2 | 0.054 (0.00 - 0.1)* | | | |
| Bai3 | 0.117 (0.015 - 0.2)* | 0.175 (0.03 - 0.371)** | | |
| Cam1 | 0.017 (-0.015 - 0.063) | 0.111 (-0.015 - 0.283) | 0.04 (-0.013 - 0.098) | |
| Isq | 0.051 (0.007 - 0.117)* | 0.137 (0.014 - 0.318)** | 0.178 (0.075 - 0.261)*** | 0.082 (0.01 - 0.159)*** |
